# Variants in RNA-Seq data show a continued mutation rate during strain preservation of *Schizophyllum commune*

**DOI:** 10.1101/201012

**Authors:** Thies Gehrmann, Jordi F. Pelkmans, Luis G. Lugones, Han A.B. Wösten, Thomas Abeel, Marcel J.T. Reinders

## Abstract

**Background:** Typical microorganism studies link genetic markers to physiological observations, like growth and survival. Experiments are carefully designed, comparing wildtype strains with knockout strains, and replications are conducted to capture biological variation. To maintain monoclonal strains, strain preservation systems are used to keep the number of generations between the primary stock and the experimental measurement low, to decrease the influence of spontaneous mutations on the experimental outcome. The impact of spontaneous mutations during the minimal number of growth cycles for the experimental design is, however, poorly studied.

**Results:** We set out to characterize the mutation landscape using a transcriptomic dataset of *Schizophyllum commune*, a laboratory model for mushroom formation. We designed a methodology to detect SNPs from the RNA-seq data, and found a mutation rate of 1.923 10^−8^ per haploid genome per base per generation, highly similar to the previously described mutation rate of *S. commune* in the wild. Our results imply that approximately 300 mutations are generated during growth of a colony on an agar plate, of which 5 would introduce stop codons. Knock-outs did not incur an increase of mutations and chromosomal recombination occurring at mating type loci was frequent. We found that missense and nonsense SNPs were selected against throughout the experiment. Also, most mutations show a low variant allele frequency and appear only in a small part of the population. Yet, we found 40 genes that gained a nonsense mutation affecting one of its annotated protein domains, and more than 400 genes having a missense mutation inside an annotated protein domain. Further, we found transcription factors, metabolic genes and cazymes having gained a mutation. Hence, the mutation landscape is wide-spread and has many functional annotations.

**Conclusions:** We have shown that spontaneous mutations accumulate in typical microorganism experiments, where one usually assumes that these do not happen. As these mutations possibly confound experiments they should be minimized as much as possible, or, at least, be trackable. Therefore, we recommend labs to ensure that biological replicates originate from different parental plates, as much as possible.

## Introduction

Experiments with microorganisms rely on the assumption that the organisms used in independent experiments are identical to ensure that differences in phenotypic characteristics are not the result of an underlying genetic heterogeneity. In reality, spontaneous mutations are regularly acquired during cell division(Baer *et al.,* 2007), invalidating this assumption. These spontaneous mutations represent confounding factors in the original experiments (Barrick and Lenski, 2013), as well as for independent replication experiments. These confounders usually go unnoticed, or are disregarded. However, a change of a specific phenotypic trait can reveal the occurrence of spontaneous mutations. For example, *Saccharomyces cerevisiae* is often plagued by the *petite* phenotype, caused by deletion of mitochondrial DNA(Zeyl and DeVisser, 2001; Joseph and Hall, 2004). As another example, the mutation rate of 2.0×10^-8 per base per haploid genome per generation(Baranova *et al.,* 2015) of *Schizophyllum commune* (model wood rot mushroom) regularly interferes with biological experiments(Raper and Miles, 1958), and consequently several mutations frequently occur in laboratory settings: the *thn* mutant prevents aerial hyphae formation(Wessels *et al.,* 1991), the *streak* mutant results in a blue color(Miles *et al.,* 1956), and the *fbf* mutation prevents mushroom formation(Springer and Wessels, 1989).

To prevent these and other mutations from seeping into other experiments, labs utilize a strain preservation system (Supplementary Note 1). In general, a strain preservation system attempts to minimize the number of generations between the primary stock and the strains from which measurements or materials are eventually sampled. This involves ensuring the long-term preservation of the primary stock of a given strain. Each lab worker creates their own personal stock, subcultured from the primary stock, and samples exclusively from this stock to perform experiments. For each experiment, a ‘mother plate’ is subcultured from the personal stock, from which all subsequent measurements are made. If a personal stock is depleted, it is recreated from the primary stock. This procedure is followed for each strain, including mutant strains derived from the primary stock. To ensure statistical rigor, experiments are replicated, often seeded simultaneously from the same mother plate to reduce human error. Although quite some mitosis steps take place in such experiments, it is presumed that there are no genetic alterations with respect to the original strain except for the intentionally introduced mutations. But, it is not clear how spontaneous mutations impact phenotypic or transcriptomic differences.

We set out to capture and characterize the mutations acquired during a typical laboratory experiment. We chose the *S. commune* mushroom because this species suffers frequently from spontaneous mutations that change the phenotype of the strain. We use a near-isogenic dikaryonic strain of *S. commune (H4-8)*, meaning that each hyphal compartment contains two nuclei. These nuclei have been backcrossed in such a way that their genomic material differs only in their mating type loci (Materials, Supplementary Note 2). As *S. commune* grows linearly outwards, with cell division occurring only at the hyphal tips, a mutation gained at any stage of this growth will naturally be passed along to its descendants.

Our dataset is composed of 46 RNA-Seq measurements (see Materials and Methods) from the dikaryotic wildtype and knockout strains of genes involved in mushroom formation (BRI1,FST3,FST4,HOM1,HOM2,GAT1,WC-1,WC-2, and C2H2). The heredity of these 46 samples is defined in the sample tree shown in Figure 1. The wild type has been sampled at five different developmental stages, across two different sequencing runs (aggregates and mushroom in the first, vegetative, vegetative induced and primordia in the second). The knockout strains were sampled at two different developmental stages (aggregates and mushroom, see Materials and Methods, Supplementary Note 3). All knockout strains are derived from a wild-type derivative, in which the ku80 gene is deactivated (node 26, Figure 1, Supplementary Note 4) to repress non-homologous chromosomal repair (De Jong *et al.,* 2010). We studied the accumulation of spontaneous genomic mutations during growth of the wildtype and knockout strains in an experiment to monitor whole genome expression during development of *S. commune* (Figure 1, Materials) (Gehrmann *et al.,* 2016; Pelkmans, 2016).

**Figure 1:**
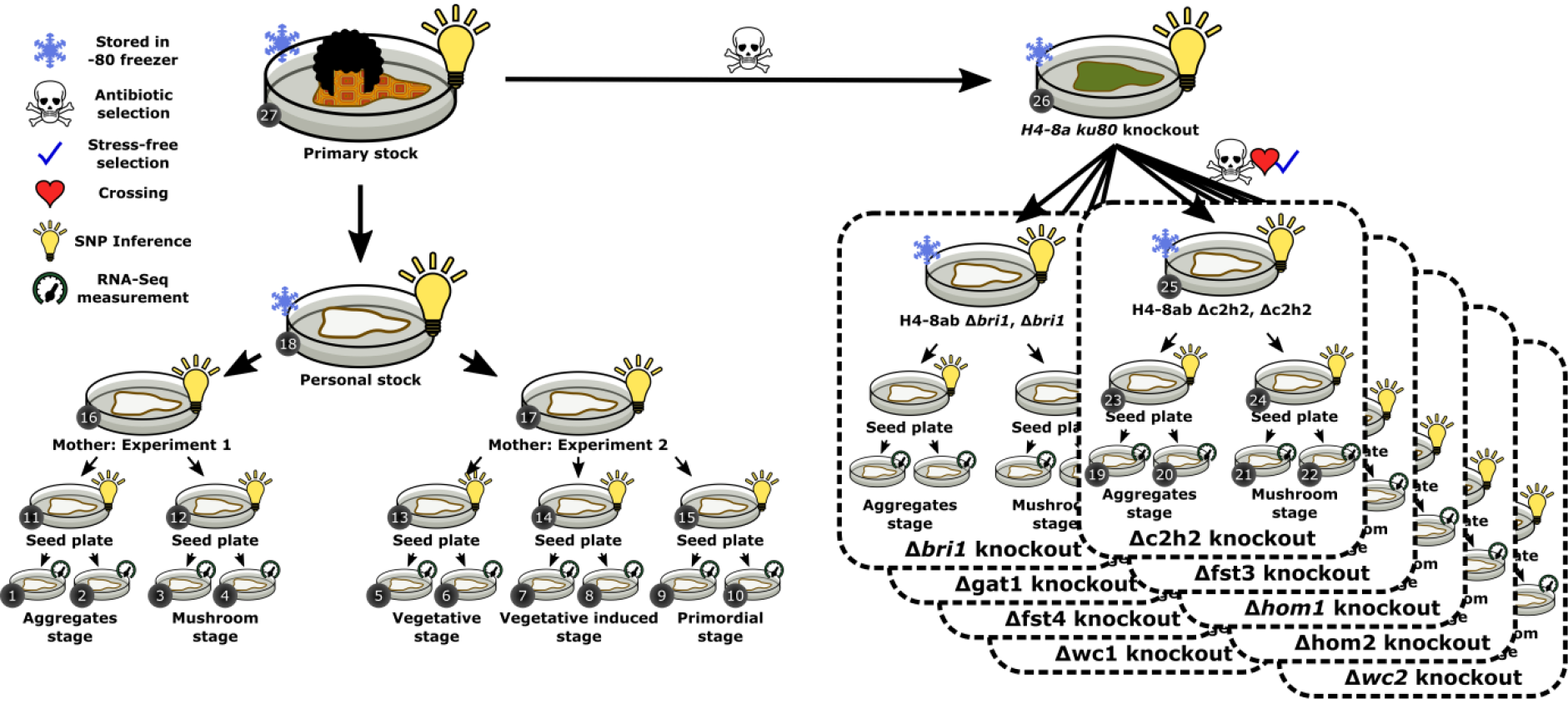
*The experimental design of our S. commune experiment. The experimental design is heavily influenced by the strain preservation system, and takes the form of a tree. The labelled nodes (1-27) are the same as those shown in Figure 2. RNA-Seq data is derived from the leaves of the tree (indicated by the measurement icon), while SNPs are inferred for the internal nodes of the tree. All nodes are samples that are originally derived from the -80 freezer (node 27), including the ku80 knockout (node 26), from which all knockout samples are derived. Initially, a researcher took a sample from the original stock, and created his own personal stock (node 18). From this, two experimental runs were conducted (nodes 16 and 18, Materials).*

## Results

### SNPs can be identified from RNA-Seq data

To characterize the mutation landscape in various steps of an experimental design, we developed a method with which single nucleotide polymorphisms (SNPs) can be identified from the transcriptome (Methods). As the genome of *S. commune* is very dense, and neighboring genes have overlapping UTRs, the transcriptome spans ~89% of the genome(Gehrmann *et al.,* 2016). This not only permits us to identify mutations that accumulated during culturing of the strain samples across a large portion of the genome but also gives us the ability to study their phenotypic effect across various growth conditions. Using the ‘*infinite sites assumption*’(Kimura, 1969), which assumes that mutations are only gained once at novel loci and never lost, we are able to associate individual SNPs to intermediate steps in our experimental setup (Figure 1). We sampled the H4-8 *S. commune* strain and nine derived knock-out strains across a total of five developmental stages (Materials). RNA was sequenced with an average coverage of 100X per sample (Materials). Using a method which leverages the lineage information in our experimental design, we identified SNPs in our RNA-Seq data (Methods).

We identified 13,249 SNPs across all our samples (Figure 1). Depending on the read depth, we detected 1,413 SNPs on average per sample (Supplementary Note 5). 94% of SNPs are heterozygous. 43 SNPs mapped to mating type regions (Materials, Supplementary Note 2). The majority (71%) of SNPs lie in gene coding regions, 27% lie in intergenic regions, and 2% lie within intron regions. We are able to capture intergenic and intronic SNPs due to the high gene density of *S. commune*, and the overlapping UTRs of transcripts. Most SNPs (92%) are present at lower abundances than the reference base, resulting in low Variant Allele Frequencies (VAF) (Supplementary Note 6). However, care should be taken to interpret the VAF, as in this case it represents a non-deconvolvable expression of subcolonal populations and nuclear specific expression. SNPs present in a large number of samples also mapped to genes that have a high RNA expression level (Supplementary Note 7). 9% of the SNPs (1,252) lie in previously predicted alternatively spliced genes (Gehrmann *et al.,* 2016).

### SNP origins reveal compounding spontaneous mutations

To investigate the accumulation of mutations throughout the experimental design, we identified the origin of SNPs in the sample tree. At each stage in the sample tree, mutations are gained with respect to the previous stage (Figure 2, see Supplementary Note 8 for the full tree). Most SNPs (85%) whose origin is identified at an internal node in the tree are supported by all its child samples (Methods), and only a small proportion (15%) are supported by a subset of child samples. As most variants match with the sample tree, it supports that the SNPs are DNA mutations rather than RNA editing substitutions. 11% (1,431) of the detected SNPs are predicted to be present in the primary stock of H4-8. The remaining 11,818 SNPs are gained at some point in the experimental procedure. Of the 783 homozygous SNPs, 85% (663) are associated to the primary stock. 7% (52) homozygous SNPs have been introduced in the ku80 knockout. The remaining 68 heterozygous SNPs are scattered throughout the experimental design, which, although unexpected due to the infinite sites assumption, may be the result of allele specific expression or silencing.

**Figure 2:**
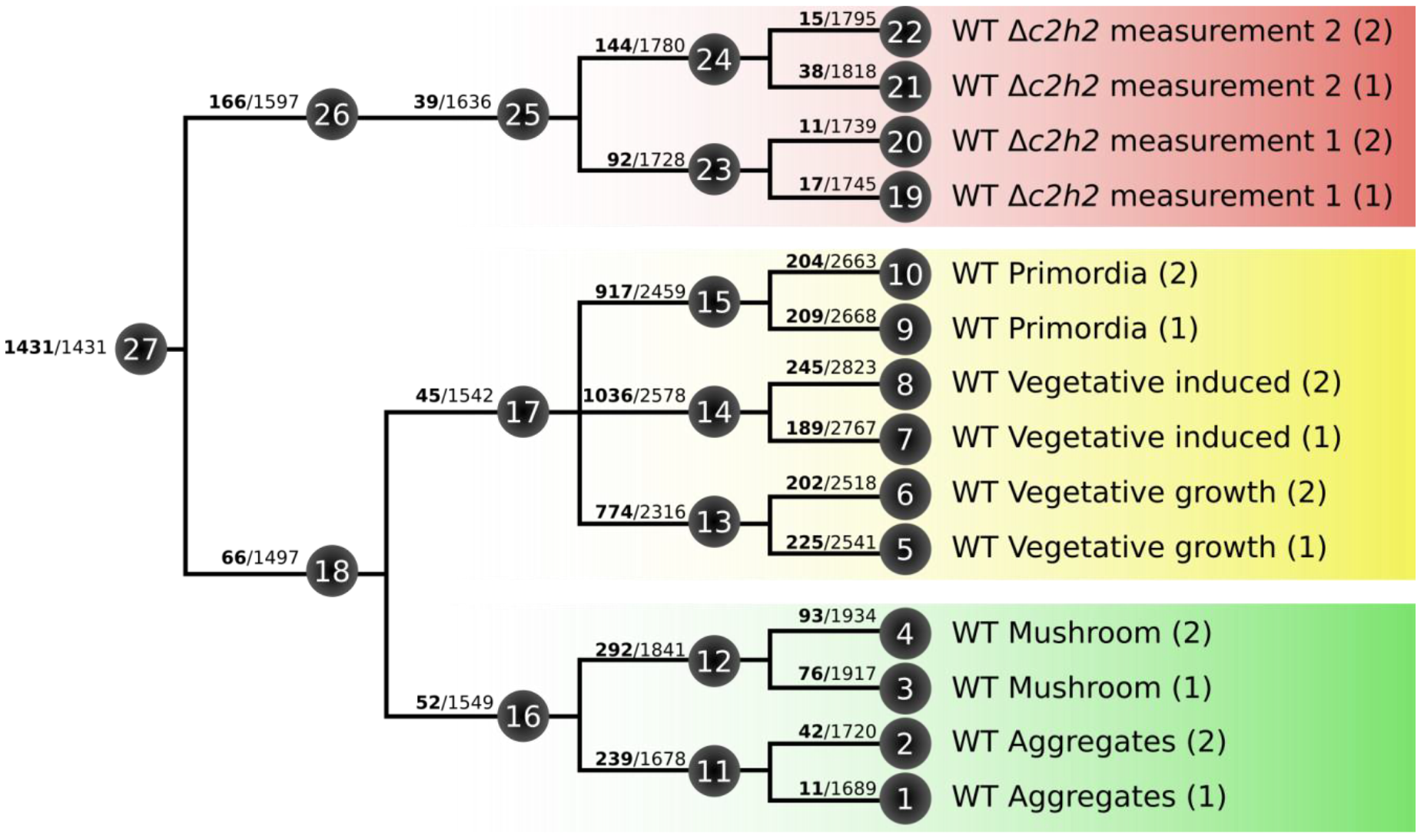
SNPs identified at each plate in the sample tree. The numbered nodes in the tree are the same as those numbered in Figure 1. Bold shows the SNPs which are gained on that plate. The nonbold number indicates the total number of SNPs observed on that sample (cumulative from root). The full tree can be seen in Supplementary Note 9.

### Estimated mutation rate is similar to the natural mutation rate

To determine if the mutation rate in the laboratory, where evolutionary stresses are removed, is different from the natural environment, we calculated a mutation rate from the observed mutations. Each plate represents approximately 200 cell divisions, and thereby 200 dikaryotic DNA duplications (Methods, Supplementary Note 9). From this, and given the 13,249 SNPs we identified, we estimated the mutation rate to be 1.9233*10^−8^ (95% CI: 1.3899*^-8^, 2.4568*10^−8^) per haploid genome per base per generation (Methods). This estimated mutation rate, in a strain preservation system, is almost identical to the mutation rate known for wild *S. commune* strains(Baranova *et al.,* 2015). The mutation rate varies across the genome (Figure 3), with 27 non-overlapping loci of 20,000bp containing more than 20 SNPs. In these regions, we observe a mutation rate of 4.158*10^−8^ (95% CI: 2.392*10^−8^, 5.924^-8^) per haploid genome per base per generation. The mating type loci exhibit a slightly (but not significantly) lower mutation rate of 1.652*10^−8^ (95% CI: 6.349×10^−9^, 2.670×10^−8^), and consequently are not enriched for or depleted of mutations (Supplementary Note 10).

**Figure 3:**
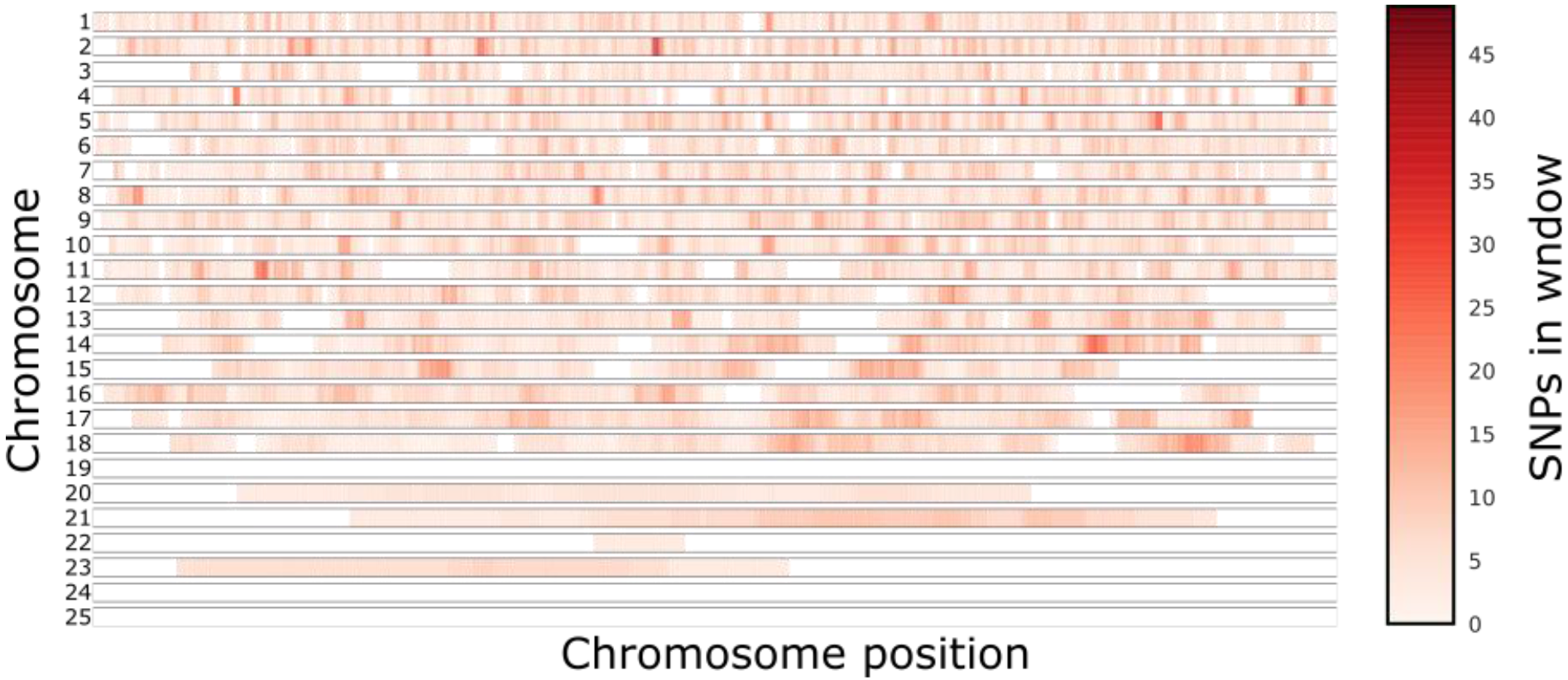
SNP hotspots across the genome. We move a sliding window across the genome (Methods), counting the SNPs in that window. The intensity of the point indicates the number of SNPs in the window around that position in the chromosome (see Methods).There are certain peaks, several of which coincide with mating type loci (chromosomes 2 and 11).

### Selection against nonsense and missense mutations

The impact of SNPs on the coding sequence was assessed. Excluding the SNPs present in the original sample, 71% (8,820) of the SNPs that were gained at some point in the experimental setup lie in coding regions (between start and stop-codons) of predicted genes. 48% (4,234) are synonymous mutations, 50% (4,769) are missense mutations, and 2% (205) are nonsense mutations. This is significantly more synonymous, less missense and less nonsense mutations than expected by random chance (p-value < 0.05, χ^2^-test) when taking into account the codon usage of *S. commune* (Supplementary Note 11). This indicates that deleterious mutations are still under negative selection in the population.

To examine the impact of these mutations on functionality, we examine five key functional groups that are highly relevant for the development and growth of this fungus. Table 1a shows that the SNPs in coding regions occur without specific enrichments across the functional groups “*transcription factors*”, “*cytochrome P450s*”, “*metabolic proteins*” and “*carbohydrate active enzymes (cazymes)*”, even for the nonsense mutations. Examining the functional groups in which these mutations lie can help us understand the impact of these SNPs on the functionality of the genes, we investigate the location of the mutation in the gene relative to the annotated protein domains. 11% (967) of the SNPs in coding regions are located within predicted domain regions of 737 genes. 47% (453) are synonymous, 50% (486) are missense, and 3% (28) are nonsense mutations. No transcription factors have mutations in their binding domains (see Table 1c). 16% (1,369) of the SNPs in coding regions are located before the end of the last annotated protein domain, also potentially altering the functional components of the protein. They lie within 1,023 genes, are annotated across various function groups (Table 1b), and are similarly distributed across synonymous 48% (661), missense 49% (668), and nonsense 3% (40) mutations as all detected SNPs. As an example, gene *G2683529* is a predicted transcription factor with a nonsense mutation upstream of its DNA binding domain. The SNP is gained in the seed plate of the first-time measurement of the Hom2 knockout. The variant has a low VAF (< 4%), and is unlikely to severely impact colony behavior due to its low expression and prevalence in the population.

**Table 1:** Number of genes with SNPs in any part of a gene in the different functional groups that lie a) anywhere in the gene, b) before the last protein domain, and c) inside a protein domain. The SNPs are split into Synonymous (S), Missense (M) and Nonsense (N) mutations. The *total* row indicates any functionally annotated gene.

|  | a) Anywhere in gene |  |  | b) Before last domain |  |  | c) In protein domain |  |  |
| --- | --- | --- | --- | --- | --- | --- | --- | --- | --- |
|  | S | M | N | S | M | N | S | M | N |
| **Transcription Factors** | 128 | 132 | 13 | 31 | 33 | 1 | 11 | 12 | 0 |
| **Cytochrome P450s** | 23 | 21 | 3 | 8 | 5 | 1 | 7 | 5 | 1 |
| **Metabolic proteins** | 177 | 183 | 13 | 55 | 77 | 2 | 47 | 71 | 2 |
| **Cazymes** | 525 | 542 | 30 | 176 | 166 | 9 | 128 | 130 | 7 |
| **Total** | **2,860** | **2,934** | **202** | **575** | **574** | **40** | **394** | **411** | **28** |

### Spontaneous SNPs may influence gene expression

Next, we set out to study the impact of detected mutations on the RNA expression level of neighboring genes. There are no SNPs within gene coding regions that do significantly influence gene expression (FDR corrected p-value > 0.05, two sample t-test with independent variance assumption between expression of samples with and without SNP). On the other hand, this is different for SNPs in (predicted) promoter regions of genes (Methods). 2% (36) of the 1,995 SNPs found in promoter regions of genes showed a significant difference in the RNA expression level of the associated genes (FDR corrected p-value < 0.01, two sample t-test with independent variance assumption between expression of samples with and without SNP). 29 of these genes have been observed to be differentially expressed between different developmental groups (Supplementary Note 12), so that the difference in expression level might not be caused by the SNP but is a result of development. For the other seven genes there was no alternative explanation. Four of them have a lower expression when the SNP is present, and three a higher. One encodes for protein ID 2539542, a (predicted) transcription factor. The SNP appeared for the first time in the aggregate stage seed plate for the *gat1* knockout and has a variant allele frequency of 11%. The observed difference between the means of the RNA expression levels is 13% with the gene having a higher RNA expression level when the SNP is present.

### During karyogamy, *S. commune* performs frequent chromosomal crossover

SNPs identified at the root node in the strain tree allow us to investigate recombination in *S. commune* (see Materials). The 1,431 SNPs predicted to be associated with the primary stock are distributed more or less evenly over the genome. Scaffold 2 and 11 show mutation hotspots (Supplementary Note 13) that map to the A and B mating type loci (Figure 4). Their exceedingly high number of mutations suggest that these regions are at the breakpoints of a chromosomal recombination. This is supported by the observation that neighboring regions are depleted of mutations, indicative of the isogenic nature of the H4-8 strain. To explain the observed recombination events at these regions, a minimum number of six chromosomal recombinations are required, amongst which one should have happened upstream, and one downstream of each mating type locus (four recombinations for the MatA loci, and two recombinations for the MatB loci). Having at least six chromosomal recombinations at these specific loci within nine backcrosses is indicative of a high chromosomal recombination frequency. We do find a few similar hotspots for the knock-out samples (Supplementary Note 14).

**Figure 4:**
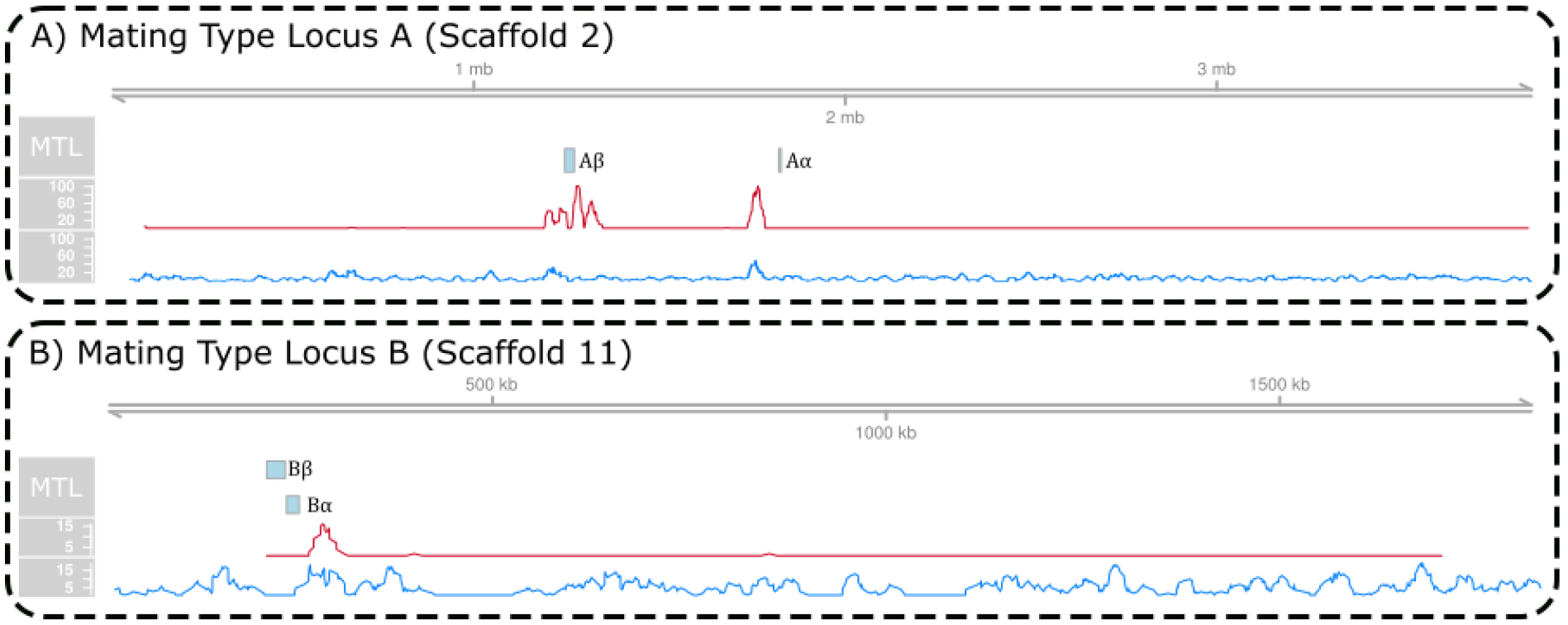
SNP hotspots around the mating type loci MatA on chromosome 2 (A) and MatB on chromosome 11 (B). The red line in each panel indicates a sliding window of 10,000bp of SNPs whose origin was at the root node of the sample tree, while the blue line indicates the SNPs whose origin was not the root node of the sample tree.

### The gene-knockout in *S. commune* procedure does not induce SNP mutations

We were wondering whether the stress conditions during the knockout procedure (see Materials) would introduce hitchhiker mutations. Between the wildtype and the *ku80* knockout, we found 166 SNPs. This is not significantly more than at any other reproductive step (node) in the tree, even after correcting for read depth (p-value > 0.05, 1 sample t-test). On the other hand, between the *ku80* knockout and the derived dikaryotic knockouts, we find significantly fewer SNPs when compared to the other reproductive steps (p-value < 0.05, 2 sample t-test with unequal variance). This indicates that the stresses of a gene knockout do not induce further spontaneous mutations.

## Discussion

The mutation rate we observe in an experimental setup in the laboratory is similar to the previously reported mutation rate identified in in wild *S. commune* strains. From this observation, it is obvious that a strain preservation system does not protect against (or lower the number of) spontaneous mutations. Consequently, even in such a laboratory setting, spontaneous mutations will confound experiments, underpinning the necessity for replication experiments. However, we do need to be careful in designing the setup of these replication experiments. If we would replicate from the same parental seed plate, similar spontaneous mutations can confound results. Hence, replications should be done from different seed plates.

We observed a similar mutation rate in the mating type loci as in the rest of the genome. James (James, 2015) argued that mushrooms have evolved a high outcrossing efficiency to ensure and encourage diversity in the population. *S. commune* has an extremely high outcrossing efficiency(Raper *et al.,* 1958). Our observation, which implies that the mating type loci are not protected from mutations, could point towards a biological mechanism that drives the high outcrossing efficiency in mushrooms. That is, the mating type loci mutations may create additional mating types. In theory, this could make it possible for hyphal anastomosis within a monokaryotic colony to result in a compatible, fertile dikaryon.

We identified a large number of SNPs originating in the primary stock (root node of the sample tree) near the mating type loci. These are indicative of recombination sites. Not having linkage information complicated the calculation of a recombination rate for *S. commune*. Based on the mutation pattern near the mating loci, we expect a high chromosomal recombination frequency for *S. commune*. Previously it has been shown that *S. commune* performs crossover at regions of high homology(Seplyarskiy *et al.,* 2014), and we see that *S. commune* also recombined very closely to mating type regions. *A. bisporus* (Pelkmans *et al.,* 2016) (for which *S. commune* serves as a model for mushroom formation) performs crossover only near the telomeric regions (Sonnenberg *et al.,* 2016). Given the seemingly alternative crossover mechanisms, we suggest caution when comparing evolutionary mechanisms between *S. commune* and *A. bisporus*.

The *ku80* knock-out strain derived from the primary stock showed a similar number of mutations as seen for the other derived strains. Initially we were expecting a higher number of mutations due to stress (e.g. passenger mutations by selection with antibiotic resistance) induced by creating the knock-out. This might not have occurred because *ku80* is involved in the DNA repair mechanism for double strand breaks. Hence, the absence of *ku80* might not have an impact on single nucleotide polymorphisms, but rather induce structural variations such as indels or inversions. Our method to detect variations of DNA from RNA sequence data was not designed to detect these larger variations and the use of short-reads for RNA-seq precludes the study of large indels and translocations that long reads would ameliorate (Salazar *et al.,* 2017). For the knock-out strains that are derived from the ku80 strain, we observed a lower mutation rate. This might be the result of crossover during the backcrossing with the primary stock wildtype to restore the ku80 gene. This would remove some mutations that were gained on one allele but not the other. The backcrossing with the primary stock wildtype might also explain the relatively high number of homozygous SNPs in the ku80 knockout strain.

As we derived mutations from RNA sequencing data, we were able to relate detected spontaneous mutations to changes in RNA expression levels (in the same samples). We found no SNPs in coding regions that influence RNA expression of the corresponding gene, and only a handful of SNPs in predicted promotor regions were associated to changes in expression levels of the corresponding genes. This suggests that regulatory regions are vulnerable to evolutionary drift, especially in intermediate plates where a large part of the functional repertoire of the organism is not utilized. This effect might even be larger, since we only used the simplistic rule of associating a regulatory SNP to a gene via its upstream promotor region. Enhancers and promotors, however, lie scattered across the genome, forming complex interactions(Vermunt *et al.,* 2014), which can be activated and deactivated by the 3D conformation of the genome(Babaei *et al.,* 2015). Detected SNPs in these regulatory regions might influence expression of a gene much further away than we now know account for. To estimate this effect, we do, however, need a more accurate picture of the complex genomic and regulatory interactions in higher fungi.

Although we did not find spontaneous mutations in coding regions to change RNA expression, we did find mutations in coding regions that led to functionally different proteins. That is, we found 411 missense SNPs in protein domains, and 40 nonsense SNPs that change the protein domain configuration of a protein. These missense and nonsense mutations are underrepresented in our observations, again pointing towards an evolutionary conservation of these regions. Nevertheless, they do involve regulatory genes and important metabolic genes. While it is known that non-essential genes evolve faster than essential genes(Jordan *et al.,* 2002), we found no functional group enriched for SNPs. As *S. commune* does not need its full functional repertoire in our experiment, we expected some groups to be more mutated than others. For example, strains used in our experiments are always grown on glucose containing minimal medium (Gehrmann *et al.,* 2016), implying that the need for carbohydrate active enzymes (*cazymes*) is reduced. Yet, we do not observe more mutations in this group of genes due to a lesser evolutionary pressure. We should, however, realize that the effect of selective pressure might be limited due to the relatively small number of generations. Based on the detected mutation rate, we can expect approximately 300 SNPs to accumulate through the growth of a single colony in a dikaryotic *S. commune* strain. Given the incidence of nonsense mutations in our dataset, we can anticipate that approximately 5 will induce a stop codon.

It is possible that a number of the SNPs we discover are the result of post-transcriptional modification, such as RNA editing. RNA-editing has been shown to occur in fungi. However, the SNPs we identify are confidently associated to nodes in an evolutionary lineage, which is not what is to be expected from RNA editing events. Additionally, the mutation rate we estimate corresponds with the known mutation rate of *S. commune* in the wild, and the SNPs around the mating type loci correctly coincide with expected recombination sites. Together, these observations indicate that the substitutions we observe are actually genetic variants, rather than post-translational modifications. To resolve these conflicting observations would require an additional study in which DNA and RNA are sampled simultaneously, such as simul-Seq(Reuter *et al.,* 2016).

It has been shown that errors in the repair of damaged DNA (and possibly cDNA) are linked to faulty variant identification(Chen *et al.,* 2016). Such errors could explain the majority of variants with low variant allele frequencies. And, in our case, the majority of SNPs do have low VAFs. However, most of our SNPs are identified across a large number of samples. Hence, it is unlikely that the DNA is damaged and incorrectly repaired at identical locations over multiple samples.

In this work, we developed an innovative method to detect SNPs in RNA-Seq data, which makes sense for *S. commune* since the transcriptome covers 89% of the genome. There have been previous attempts to call SNPs from RNA-Seq reads(Ramirez-Gonzalez *et al.,* 2015; Deelen *et al.,* 2015; Piskol *et al.,* 2013; Quinn *et al.,* 2013; Piechotta *et al.,* 2017). With the exception of JACUSA(Piechotta *et al.,* 2017), which was designed for the identification of RNA editing events, other approaches generally rely on GATK(McKenna *et al.,* 2010), which was primarily not designed for the study of variants in RNA-Seq data. Most importantly, GATK assumes an approximately uniform distribution of reads across the genome, which is certainly not the case for RNA-seq data. Furthermore, the allelic imbalance due to allele specific expression (or, in our case, karyollele specific expression) severely hampers the performance. The best practices as described by Broad Institute indicate that results are only acceptable when strict filters are used (https://software.broadinstitute.org/gatk/documentation/article.php?id=3891). When we applied the GATK pipeline to our data, we found only 351 SNPs that associated to our experimental design tree. Therefore, we chose to develop our own method. Our initial SNP calling step is permissive and will call many spurious SNPs. Our method, therefore, strongly relies on a second step to filter spurious SNPs based on prior knowledge of the evolutionary relationship of the samples. It only permits SNPs with low RNA-seq coverage if they are present across several related samples. Without knowing the relationship between the samples, this becomes very difficult.

As we derived mutations from RNA-Seq data, we do have to make a note of caution on our findings as they depend on the expression level of a gene. That is, when a gene is not expressed, no mutation can be detected. We remedy this by exploiting the (full) experimental setup. Throughout the complete experiment, only 1,612 (9.8%) of all predicted genes were considered to be not expressed (FPKM < 1) in any sample. Thus, although we do not capture the entire genome, we capture a considerable portion of it, and the reported mutation rate takes this into account.

## Conclusion

In the laboratory, the selection pressures that shape the genotype and phenotype of wildtype organisms are replaced, relaxed, or even lost. We have shown that *S. commune,* a model organism for mushroom formation, has the same high mutation rate in the lab as in the wild. Spontaneous mutations will accumulate in experiments and tven the best strain preservation system cannot prevent this. We showed that SNPs are introduced in a variety of important functional groups, and that they can have an effect on the function and regulation of genes. It is not clear that there is a better way to prevent the accumulation of spontaneous mutations, other than reducing the number of generations between the primary stock and the experimental strains derived thereof. We recommend that labs implement a sample tracking system in the lab, whereby each sample that enters a freezer is registered with its ancestor sample. This will enable the isolation of mutations should they later be discovered. Additionally, the experimental design should take into account the additional mutations that could accumulate, and replicates should originate from different parental plates. Although this may result in higher biological variation between the replicates, it will eliminate differences that result from confounding mutations that accumulated in the tree.

## Materials and methods

### H4-8 S. commune strain

The H4-8 *S. commune* strain(Ohm, de Jong, Lugones, *et al.,* 2010) is a co-isogenic dikaryon, meaning that it is a heterokaryon whose constituent homokaryons are supposedly identical with the exception of the mating type loci. It is the result of an integration of the H4-8a and H4-8b homokaryons. The H4-8b strain was achieved through 9 backcrosses between H4-8a and 4.40, selecting in each stage for a crossing that had a compatible mating type to H4-8a. During meiosis, the chromosomes are exchanged and (often) undergo crossover at locations of genetic similarity(Seplyarskiy *et al.,* 2014). The exact efficiency of this backcrossing procedure in terms of homozygosity, especially in the chromosomes containing the mating type loci is unknown.

### Mating type loci

The two homokaryons of H4-8 differ in their A and B mating type loci(Ohm, de Jong, Lugones, *et al.,* 2010). These loci were identified in version 3 of the H4-8 genome by mapping the genes annotated in the matAα, matAβ, matBα and matBβ of version 2 to the version 3 genome using the BLAST functionality of the JGI DOE website. See Supplementary Note 2.

### Knock-out strains

The knockout strains all originate from a *ku80* knockout(Ohm, de Jong, Berends, *et al.,* 2010) (Supplementary Note 4), which was used to generate a series of regulatory gene knockouts(Pelkmans, 2016), all stored in the −80 freezer. The *ku80* knockout is the result of several stressful interventions (Supplementary Note 4), over an unknown number of generations. Beyond the phenotypic and transcriptomic differences induced by the knockouts (Ohm *et al.,* 2011; Pelkmans, 2016), it is not known what additional sequence variation is induced by the knockout of the *ku80* gene and the final regulatory genes. After the knockout of the second gene, the *ku80* gene is crossed back into the genome.

### RNA-Seq data

RNA-Seq samples were retrieved from BioProject PRJNA323434. To produce these samples, mRNA was isolated from *S. commune* strain H4-8 grown at 25°C on minimal medium containing 1% glucose and 1.5% agar(Van Peer *et al.,* 2009). The wildtype strain was initially sampled twice, once in the aggregates stage of development, and once in fruiting body of the mushroom and sequenced on an Illumina Hi-Seq 2000. Knockout strains were sampled when they reached the aggregates state and mushroom stage. If a knockout was halted in an earlier developmental stage (Supplementary Note 3), then they were sampled when the wildtype reached the aggregates or mushroom stage. A later second sequencing run sequenced wildtype samples at three additional developmental stages, vegetative growth, induced vegetative growth (after exposure to O2 and light), and primordia. Details on the sequencing runs can be found in Supplementary Note 15.

### Read Alignment

Raw reads were trimmed using TRIMMOMATIC(Bolger *et al.,* 2014) and the resulting reads were aligned to the reference genome using two-pass STAR(Dobin *et al.,* 2013), where the second pass used the splice junctions detected in all samples during the first alignment pass (Supplementary Note 16). Reads that ambiguously mapped were discarded. STAR was used to sort the resulting BAM files based on read alignment co-ordinate. Duplicate reads were flagged with PICARD (http://picard.sourceforge.net./).

### Detecting SNPs from RNA-Seq data

We process the aligned BAM files, ignoring duplicate reads, counting the number of observed nucleotides at each position in the genome. If the quality of a base is less than 30 in the PHRED scoring system, it is not counted. Based on the CIGAR strings in the BAM file, we also maintain a record of insertions and deletions observed in the alignments. For each position on the genome, we test a base for SNPs only if the base is not within 4 bases of a possible insertion/deletion site/splice junction, and that base is not an ‘N’ in the reference genome. For each nucleotide, we calculate the probability that it is observed erroneously. To do this, we assume that each erroneous observation of a nucleotide at a specific locus follows a Bernouilli trial with a small probability of success. With multiple observations, we build a binomial distribution around the probability of observing a specific nucleotide by error. Thus, when a locus has a depth of *x*, and x_*n*_ counts of nucleotide *n*, then P(*X*_*n*_ > *x*_*n*_) expresses the probability of observing more than *x*_*n*_ counts erroneously, where *X*_*n*_ ~ B(*x*_*n*_, 0.01). Clearly, with increasing observations of the nucleotide (*x*_*n*_), the probability of seeing that nucleotide at that locus as the result of an error becomes smaller. If this probability becomes smaller than 0.05 we conclude that the nucleotide is not observed erroneously, and thus is truly observed. We do so for all four nucleotides and when one of them passes this test and it is not equal to the reference nucleotide of that locus, we call a potential SNP at that base. Any SNP in a gene knockout region that originates from that knockout sample is removed. All positive and negative base calls are output in VCF format.

### Assigning SNP origin in the generation tree

We use the lineage information in the sample tree to enhance our confidence in a SNP, and to remove spurious SNP calls. The VCF files of all the samples are merged and sorted on base coordinates. SNP calls from different samples are grouped together at each base. A generation tree is constructed, such as the one shown in Figure 1. For each SNP, we determine which nodes in the tree are possible candidates for the origin of the SNP. To do this, we calculate three metrics for each node of the tree: *S*_*n*_, the number of descendent leaf nodes that have this SNP; *E*_*n*_, the number of reads supporting this SNP across all the child nodes, and *P*_*n*_, the *possibility* of this node harboring the SNP, being either ‘*yes*’, ‘*no*’ or '*maybe*’. For leaf-nodes a SNP is ‘yes’ when the SNP is present, ‘no’ when the SNP is not present, and ‘maybe’ when there is not enough depth to make a SNP call (i.e. depth < 3). For non-leaf nodes, the possibility of a SNP is ‘yes' when all its descendent leaves are ‘*yes*’, ‘*no*’ when at least one of its descendent leaves is ‘*no*’, and ‘*maybe*’ when all its descendent leaves are ‘*yes*' or ‘*maybe*’. For each SNP, we select the nodes highest (towards the root) in the tree where *P*_*n*_ is ‘yes’ or ‘maybe’, and either *E*_*n*_ > 3 or *S*_*n*_ > 1, as the node of origin for that SNP. This results in SNPs that are either supported by sufficient depth within at least one sample, or supported by multiple samples. SNPs with multiple alternative nucleotides, or SNPs whose origin can’t be resolved (i.e. no origin found, or multiple origins found) are discarded.

### Estimating transcript abundance

To calculate transcript abundance, we pre-processed the reads with TRIMMOMATIC(Bolger *et al.,* 2014) and aligned the reads to the genome using a two-pass STAR, as in the read alignment above, only in this case we permitted ambiguous alignments. Expression of each transcript for each sample was quantified and normalized with the Cufflinks(Trapnell *et al.,* 2012) toolkit.

### Associating SNPs to genes and assessing deleteriousness

If a SNP lies within the coding region of a gene, then we can assess the deleteriousness of the SNP. If the transcript with the SNP produces the same amino acid sequence as without, then the SNP is considered synonymous. If, on the other hand, the amino acid sequence is changed, then it is a missense mutation, and if the sequence is shortened, then it is described as nonsense. If a SNP lies within 500bp upstream of the start codon of a gene, then we say that the SNP falls within the promotor region of that gene.

### Calculating a mutation rate

To calculate a mutation rate, we consider the number of mutations associated to each node in the tree. As we do not precisely know how many steps were involved in creating the original double knockouts plates, the ku80 knockout plate, and the primary stock plate, we excluded those samples from the calculation. The number of SNPs in each sample are divided by the number of bases considered, and multiplied by the number of generations that each plate represents (200, see Supplementary Note 9 and (Raudaskoski and Salonen, 1984)). The mutation rates for each plate are averaged to arrive at a cross-sample mutation rate. A confidence interval is calculated assuming a normal distribution. For the genome-wide mutation rate, we used the number of bases covered with at least 5 reads (excluding mating type loci) multiplied by two, representing the callable part of the diploid genome. Because the number of detected mutations is dependent upon read depth, we need to correct for the library sizes in each sample. However, since the ability to call a SNP depends on the coverage of each base, we should, more specifically, correct for the number of confidently callable bases per sample. For each sample *i*, *k*(*i*) denotes the number of bases with at least 5x coverage (Supplementary Note 15). For non-leaf nodes, we infer *k*(*i*) to be the maximum across its leaves. The number of SNPs detected in each sample is multiplied by 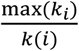 where *max*(*k*_*i*_) is the maximum number of confidently callable bases across all the samples. This scales the number of mutations up for those samples with less read depth.

### Detecting mutation hotspots

A sliding window of 10,000bp up- and down-stream of a detected SNP, which contains at least 20 SNPs is considered a mutation hotspot.

### Functional annotations

Interpro domain annotations were taken from the JGI DOE website, filtered with a score threshold < 0.05. Transcription factors were predicted based on a curated list of fungal DNA binding domains, as in (Gehrmann *et al.,* 2016). Cytochrome P450 genes were predicted based on the Interpro domain IPR001128, and metabolic genes based on the GO annotation GO:0008152. Carbohydrate active proteins were predicted using the CAT(Park *et al.,* 2010; Lombard *et al.,* 2014) tool, selecting only those proteins which are predicted both with the PFAM and sequence predictors. Alternatively spliced genes were taken from (Gehrmann *et al.,* 2016).

## Acknowledgements

The sequence and annotation data of *S. commune* H4-8 version 3 were produced by the US Department of Energy Joint Genome Institute http://www.jgi.doe.gov/ in collaboration with the user community. This research is supported by the Dutch Technology Foundation STW, which is part of the Netherlands Organisation for Scientific Research (NWO), and which is partly funded by the Ministry of Economic Affairs. Additionally, we thank Aurin M. Vos and Robin A Ohm for fruitful discussions.

### Author contributions

TG, HABW, TA, and MJTR wrote the manuscript. JFP and LGL performed the biological experiments. TG, TA, and MJTR designed the analyses. TG performed the analyses. All authors aided in biological interpretation of the results. All authors reviewed the manuscript.

